# Capillary hemoglobin electrophoresis of healthy and anemic dogs: quantification, validation, and reference intervals of hemoglobin fractions

**DOI:** 10.1101/633198

**Authors:** Ioannis L. Oikonomidis, Theodora K. Tsouloufi, Mathios E. Mylonakis, Maria Kritsepi-Konstantinou

## Abstract

Despite the advances in canine medicine and the rapid gaining of attention of canine models in biomedical field and particularly in hemoglobin genes research, the studies on canine hemoglobin composition are sparse with ambiguous findings. Our aim was: i) to investigate the electrophoretic pattern of canine hemoglobin and the possible effect of age, sex, and anemia using a capillary electrophoresis assay, and ii) to validate this assay and calculate reference intervals (RIs) for canine hemoglobin fractions. Blood samples were collected from 53 healthy and 42 dogs with regenerative and non-regenerative anemias. The Sebia Capillarys 2 flex-piercing was used for hemoglobin analysis and it was validated using canine blood samples. R statistical language was employed for the statistical analyses. A major hemoglobin fraction (named HbA_0_) and a minor one (named HbA_2_) were identified in 100% and 47.4% of samples, respectively. The within-run and between-run CV was 0.1% for HbA_0_ and 9.1% and 11.2% for HbA_2_, respectively. The extremely narrow range of HbA_0_ and HbA_2_ values hampered a linearity study using canine blood samples. The RIs for HbA_0_ and HbA_2_ were 98.9-100% and 0-1.1%, respectively. HbA_0_ and HbA_2_ values were not correlated with age (*P*=0.866). No differences were observed in the median HbA_0_ and HbA_2_ between the two sexes (*P*=0.823), and healthy and anemic dogs (*P*=0.805). In conclusion, the capillary electrophoresis revealed a major hemoglobin fraction and an inconsistently present minor fraction. No effect of age, sex, or anemia was detected. The assay used was validated and RIs were generated, so as to be suitable for use in future investigations.

## Introduction

Hemoglobin is the oxygen-carrying moiety of erythrocytes. Structurally, it is a globular polypeptide tetramer, which consists of two pairs of unlike globin chains that form a shell around a central cavity. The latter contains four oxygen-binding heme groups, each of which is covalently linked to a globin chain.

In healthy humans, hemoglobin consists of: i) a major fraction, HbA_0_ (α_2_β_2_), which comprises approximately 95% of the total hemoglobin; ii) a minor fraction, HbA_2_ (α_2_δ_2_), which is normally less than 3.5% of total hemoglobin and iii) the fetal hemoglobin, HbF (α_2_γ_2_) [1]. In human medicine, more than 700 hemoglobinopathies have been described to date with most of them being clinically benign [2]. The term hemoglobinopathy is broadly used to describe both quantitative (thalassemias) and qualitative (true hemoglobinopathies) hemoglobin disorders [3]. However, in a strict sense, hemoglobinopathies and thalassemias are two genetically distinct groups of diseases, although clinical manifestations may overlap [1]. Specifically, thalassemias are characterized by a reduced production of the normal globin chain and may result from gene deletion or mutations that affect the transcription or stability of mRNAs [1]. On the other hand, the vast majority of hemoglobinopathies, including the clinically important ones, result from single nucleotide substitutions that are translated to single amino acid substitutions, primarily in the non-α chain, causing alterations in the secondary and tertiary structures of hemoglobin tetramer [1, 4].

High pressure liquid chromatography (HPLC) and capillary zone electrophoresis (CZE) are the most widely used methods for human hemoglobin analysis and for the initial diagnosis of hemoglobinopathies, both of which have superior analytic and diagnostic performance when compared to other available methods, such as gel electrophoresis and mass spectroscopy [5]. CZE allows the successful separation of the normal human hemoglobin fractions, but it can also detect abnormal hemoglobin variants with altered charge resulting either from mutations that directly influence the charge of the molecule or indirectly from mutations that alter the higher-order structure [4]. In particular, Sebia Capillarys 2-flex piercing (Sebia, Norcross, USA), the updated model of Sebia Capillarys, has been successfully validated for human hemoglobin analysis and diagnosis of hemoglobinopathies [6]. Additionally, the same analyzer has been recently successfully validated for the measurement of the major fraction of glycated hemoglobin (HbA_1c_) in dogs [7].

Currently, there is a dearth of published studies on hemoglobin composition in dogs and they have been conducted almost half a century ago [8, 9], although dogs are rapidly gaining attention as potential models in various biomedical areas, while they are considered the ideal model particularly for the study of hemoglobin genes [10]. According to the above cited studies, no HbF is recognized in dogs, while a minor hemoglobin fraction may be detected [8, 9]. However, no further information is provided about the prevalence, quantification, and electrophoretic characteristics of the minor hemoglobin fraction. Only recently the minor hemoglobin fraction was quantified using acetate cellulose electrophoresis [11]. Surprisingly, the authors of this study also reported the presence of HbF in adult dogs, raising questions about our prior knowledge, but also about the utility of different assays for canine hemoglobin analysis [11].

In the aforementioned context, the objectives of this study were: i) to investigate the electrophoretic patterns of canine hemoglobin using a new automated capillary electrophoresis assay; ii) to study the effect of age, sex, and anemia unrelated to hemoglobin disorders on the electrophoretic pattern of canine hemoglobin; and iii) to validate the herein used assay for canine hemoglobin analysis and calculate appropriate reference intervals, so as to be suitable for use in future studies or in the clinical setting.

## Materials and methods

The blood samples used in this study were aliquots of specimens collected (owners’ consent provided) for diagnostic purposes, routine health check, or pre-operatively from healthy dogs referred to the Companion Animal Clinic, School of Veterinary Medicine, Faculty of Health Sciences, Aristotle University of Thessaloniki, Greece. The reference individuals were selected by a direct a priori method, based on the following inclusion criteria: age >6 months, up-to-date vaccination and deworming status, no history of illness or medication in the preceding month, unremarkable physical examination, and normal complete blood count. Blood samplings were performed at admission by jugular venipuncture and the samples were collected into K3-ethylene diamine tetra-acetic acid (EDTA) coated tubes (Deltalab, Barcelona, Spain). Anemia was defined as red blood cell count <5.36 × 10^9^/L, or hemoglobin concentration <122 g/L, or hematocrit <0.372 L/L [12]. The anemia was classified as regenerative when the absolute reticulocyte count was >60,000/μL [13]. Grossly hemolysed (in vitro hemolysis) and lipemic samples were excluded from the study. A complete blood count was performed on the Advia 120 hematology analyzer (Siemens Healthcare Diagnostics, Deerfield, USA) within 2 h of sampling.

Hemoglobin electrophoresis was carried out within 4 h of sampling. Routine maintenance, assay and internal quality control procedures were conducted as defined in the analyser manuals. A normal electrophoretogram from a human patient was used for comparison. The automated analyser, Sebia Capillarys 2 flex-piercing, and the dedicated kit (Sebia, Norcross, USA) were used for the detection and quantification of different canine hemoglobin fractions as a percentage of total hemoglobin. The principle of the Capillarys 2 flex-piercing assay is CZE, in which charged molecules are discriminated by their electrophoretic mobility in an alkaline buffer (pH 9.4). The analyser is equipped with eight silica capillaries, which enable the simultaneous analysis of eight whole blood samples. In brief, the EDTA-treated whole blood sample is diluted with a hemolysing solution and the resulting solution is then hydrodynamically injected at the anodic end of the capillary. A constant, high voltage is applied for 8 min, which allows the migration and separation of the hemoglobin variants. These are then directly detected by spectrophotometry (415 nm) and the electrophoretograms are automatically generated. The total output time is approximately 20 min for the first run and 12 min for every other run.

The validation of the analyzer was initially designed to include linearity, repeatability, and reproducibility. The repeatability or within-run precision was evaluated using blood samples from three dogs. Each sample was measured eight times in succession and the coefficient of variation (CV) was calculated. Blood samples from the same three dogs were used for the evaluation of reproducibility or between-run precision. Six aliquots were made from each sample and were measured over a period of 3 days; then, the CV was calculated.

The distribution of data was assessed using the Shapiro–Wilk test. The 95% reference intervals (RIs) were calculated using the non-parametric method, while the 90% confidence intervals (CIs) for the lower and upper reference limits were calculated by the bootstrap method. Cook’s method was employed for the detection of outliers. For the determination of reference intervals, the R package referenceIntervals was used. The exact Wilcoxon and Kruskal-Wallis rank-sum tests were employed for median comparison between two or three different groups, respectively. Spearman’s rank correlation coefficients were used for correlation analyses. All the statistical analyses were conducted using the statistical language R (R Foundation for Statistical Computing, Vienna, Austria). Level of significance was set at 0.05 (P<0.05).

## Results

In total, 95 dogs were sampled. The reference population comprised 53 dogs (27 males and 26 females) with mean (±SD) age of 6.0±3.8 years and hemoglobin concentration of 155±16 g/L. The anemic population comprised 42 dogs (19 males and 23 females) with mean (±SD) age of 6.6±4.1 years and hemoglobin concentration of 75±27 g/L. The anemia was classified as non-regenerative in 16/42 (38.1%) dogs and regenerative in 26/42 (61.9%) dogs.

The inspection of the electrophoretograms revealed one major and one minor hemoglobin fraction. The major canine hemoglobin fraction migrated slower towards the anode than the respective human HbA_0_ (Fig 1) and it was consistently present in all examined samples (95/95, 100%). The minor fraction migrated slightly slower towards the anode compared to human HbA_2_ and it was evident in 26/53 (49.1%) reference individuals and in 19/42 (45.2%) anemic dogs. For the purposes of this study, we refer to the major canine hemoglobin fraction as HbA_0_ and to the minor one as HbA_2_.

**Fig 1.**
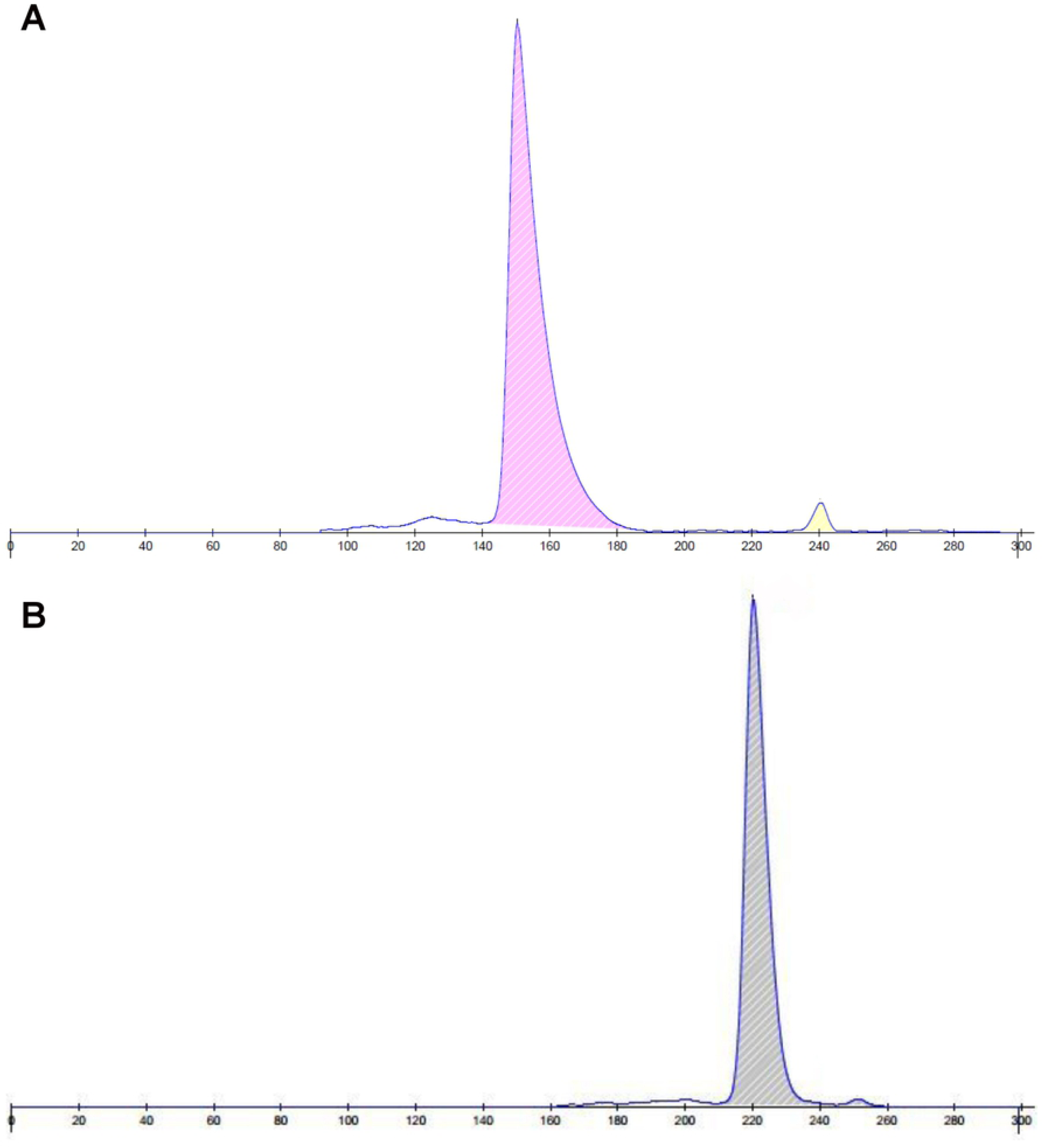
Two representative hemoglobin electrophoretograms from a healthy human (A) and a healthy dog (B). The major (HbA0) and the adult minor (HbA2) hemoglobin fractions are depicted in both electrophoretograms. The major canine hemoglobin fraction migrates slower towards the anode than the respective human one. The minor fraction migrates slightly slower towards the anode compared to human HbA2 and it is inconsistently present in dogs.

The total within-run and between-run CV for HbA_0_ was 0.1%, while for HbA_2_ was 9.1% and 11.2%, respectively. Specificity (dilutional linearity study) using canine blood samples could not be performed due to the extremely narrow range of HbA_0_ and HbA_2_ percentages in our canine population. No outliers were detected in the reference population using Cook’s method. The 95% RI for HbA_0_ was 98.9-100% with the CIs for the lower and upper reference limits being 98.8-99.0% and 100%, respectively. The 95% RI for HbA_2_ was 0-1.1% with the CIs for the lower and upper reference limits being 0 and 1.0-1.2%, respectively.

HbA_0_ and HbA_2_ values were not significantly correlated with age (*P*=0.866). No statistically significant difference (*P*=0.823) was observed in the median HbA_0_ and HbA_2_ between male and female dogs. The median (range) HbA_0_ and HbA_2_ was 100% (98.9-100%) and 0% (0-1.1%), respectively, in both sexes. No statistically significant difference (*P*=0.805) was detected in the median (range) HbA_0_ and HbA_2_ between the reference population [100% (98.9-100%) and 0% (0-1.1%), respectively] and dogs with non-regenerative [100% (98.9-100%) and 0% (0-1.1%), respectively] or regenerative anemia [100% (99.0-100%) and 0% (0-1.0%), respectively] (Fig 2).

**Fig 2.**
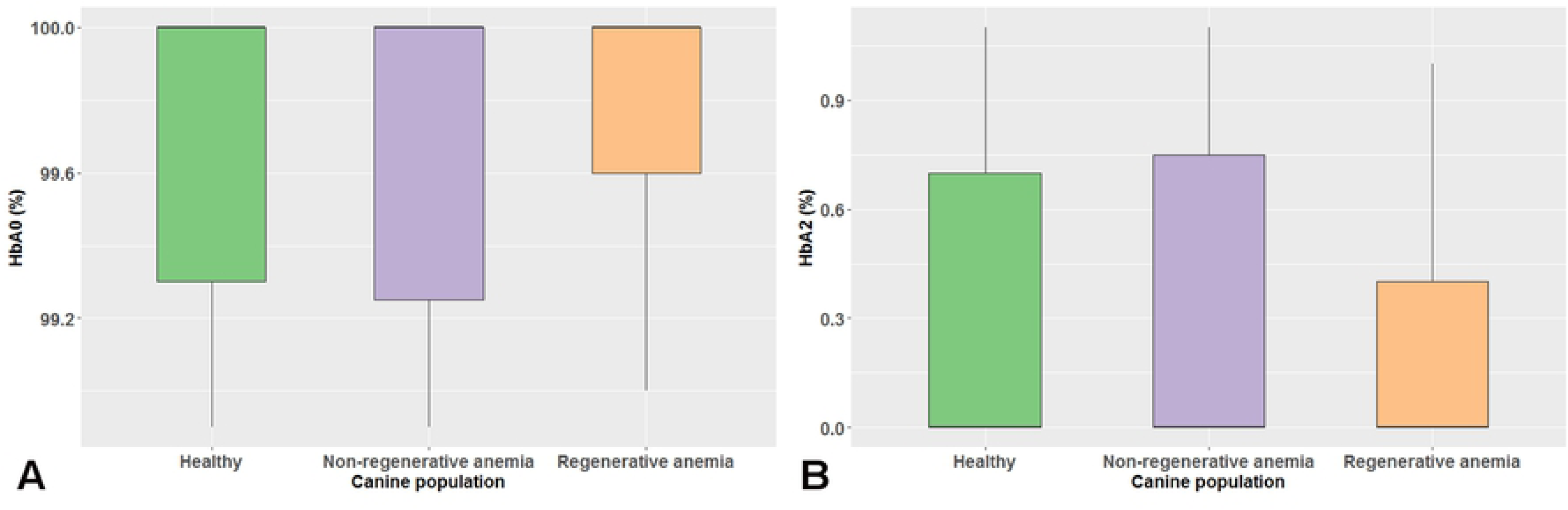
Boxplots of the major (A) and minor (B) hemoglobin fraction values of the reference populations and dogs with non-regenerative or regenerative anemia are depicted. The colored boxes represent the main body of data; they are bisected by a line, which stands for the median value. No statistically significant difference (*P*=0.805) was detected in the median values of both canine hemoglobin fractions between the three groups.

## Discussion

In this study, the electrophoretic pattern of canine hemoglobin was investigated using a new automated capillary electrophoresis assay. This assay was validated for canine hemoglobin analysis and appropriate reference intervals were calculated for adult dogs. The effect of age and sex on canine hemoglobin electrophoretic pattern was also evaluated. Finally, we investigated if anemias that were not attributed to a hemoglobin disorder, could affect the hemoglobin electrophoretic pattern.

The inspection of the electrophoretograms revealed two hemoglobin fractions: one major fraction that was constantly present in all of the enrolled dogs and one minor fraction that was detected in approximately half of the dogs. The major canine hemoglobin fraction was found to migrate slower towards the anode compared to human HbA_0_, while the minor canine hemoglobin fraction migrated slightly slower than human HbA_2_. A third hemoglobin fraction consistent with HbF was not detected in any of the dogs included in this study. Our findings are in agreement with previous studies using gel electrophoresis, which reported the absence of HbF and the presence of one or two hemoglobin fractions in dogs [8, 9]. However, in the aforementioned studies, no further information was provided about the electrophoretic features, the prevalence, and the quantification of the different hemoglobin fractions. The characterization of HbA_2_ was only recently done in canine samples [11]. In this study, the prevalence of HbA_2_ in healthy dogs was higher, yet similar to ours (64.1% versus 49.1%, respectively). However, the range of HbA_2_ value was wider and roughly three times the one reported in our study. However, Atyabi et al. surprisingly reported the presence of HbF in 50.0% of their samples [11], as opposed to current and previously published studies [8, 9], which reported the absence of canine HbF. The source of the observed discrepancy between the study of Atyabi et al. [11] and the rest of the published studies, including the present one, cannot be easily explained. Be that as it may, both preanalytic (handling and storage of the blood samples) and analytic factors (inherent limitations of the used method) may have contributed to the observed differences. This further underlines the need to utilize contemporary methods and properly validate them for use in different species.

HPLC and CZE are the most widely used methods for human hemoglobin analysis and for the initial diagnosis of hemoglobinopathies [5]. These two methods have comparable results and share some major advantages, such as the accuracy, rapidness, and high throughput; however, each of them has its disadvantages, primarily referring to inability for identification of some human-specific hemoglobin variants [5]. However, a major advantage of CZE over HPLC, which is potentially applicable to different species, is the substantially better visualization of the results; indeed, post-translational modification and degradation peaks are often present in HPLC chromatograms, potentially making the interpretation problematic [5]. Gel electrophoresis and mass spectroscopy can likewise be used for the hemoglobin analysis and diagnosis of hemoglobinopathies; notwithstanding, a major disadvantage is recognized in both of them. Gel electrophoresis is characterized by an inherent lower accuracy and sensitivity [5], while mass spectroscopy is unable to detect intact globin chains with a slightly different mass, reportedly less than 6 Da [14].

The capillary electrophoresis assay used in this study has been recently successfully validated for the measurement of canine HbA_1c_ [7]. However, to our knowledge, this is the first time that this assay is utilized for canine hemoglobin electrophoresis and thus, a study of the analytic performance of this assay is valuable. The repeatability and reproducibility of this assay for HbA_0_ measurement, using canine blood samples, was excellent and in agreement with studies in human medicine [6]. However, the within-run and between-run CV for HbA_2_ measurement was considerably higher than the one reported for human HbA_2_ [6]. The higher imprecision in canine HbA_2_ measurement can be attributed, at least partially, to the extremely low values of the HbA_2_ in dogs, which are not normally seen in humans; however, the performance is likely acceptable for use, although this cannot be clearly stated given the absence of specific performance goals in dogs. It should be noted that none of the previously used assays for canine hemoglobin electrophoresis was validated for use in dogs. Additionally, appropriate RIs were calculated for adult dogs with the range for both hemoglobin fractions being narrower compared to human one. Finally, the age and sex do not appear to have an effect on canine hemoglobin electrophoretic pattern, in accordance to human studies reporting only a minimal, effect of age and sex, and the study by Atyabi et al. which found no difference between male and female dogs [11, 15].

Given that anemia (of variable severity) is the usual clinical manifestation of hemoglobinopathies in humans [3], we also decided to investigate whether anemias (regenerative or non-regenerative) that were not related to hemoglobin disorders, might have an effect on the electrophoretic pattern of canine hemoglobin. No quantitative or qualitative hemoglobin abnormalities were detected in the electrophoretic pattern of anemic dogs when compared to our reference population. In spite of the small sample size of anemic dogs, this finding indicates that an anemia not attributable to a hemoglobin disorder does not interfere with the capillary electrophoresis assay used in our study.

## Conclusions

The canine hemoglobin consists of a major fraction and a minor one, inconsistently present in very low proportions. A new automated capillary electrophoresis assay was validated for the separation of canine hemoglobin fractions and appropriate RIs were generated. Our study indicates no age or sex effect on hemoglobin electrophoretic pattern among adult dogs, while no quantitative or qualitative hemoglobin abnormalities were detected in the anemic dogs without evidence for a hemoglobin disorder. The capillary electrophoresis assay used in this study is the only validated assay that can be used in future research studies on canine hemoglobin or in clinical cases suspected of having a hemoglobin disorder.

## Acknowledgments

The authors have no aknowledgements to state.

